# Substantial contribution of in-situ produced bacterial lipids to the sedimentary lipidome

**DOI:** 10.1101/2025.02.27.640310

**Authors:** Su Ding, Nicole J. Bale, Anna Cutmore, F. A. Bastiaan von Meijenfeldt, Stefan Schouten, Jaap S. Sinninghe Damsté

**Affiliations:** NIOZ Royal Netherlands Institute for Sea Research, Department of Marine Microbiology and Biogeochemistry, Texel, The Netherlands; Department of Earth Sciences, Faculty of Geosciences, Utrecht University, Utrecht, The Netherlands

## Abstract

The sedimentary lipid pool is comprised of a myriad of individual components. Due to their importance for organic carbon sequestration and their application in paleoclimatic and geobiological reconstructions, its composition has been studied for many decades with targeted approaches but an overall view on its composition is still lacking. In part this uncertainty relates to the different sources of sedimentary lipids, they can be both delivered from the overlying water column by sedimentation, but also produced in-situ by sediment dwelling organisms. Another uncertainty relates to the differing degree of preservation, both between lipid groups and relative to other organic matters. Here we conduct an untargeted analysis of the sedimentary lipidome in the Black Sea using high resolution mass spectrometry. Besides commonly reported phytoplankton-derived fossil lipids, a diverse and abundant set of sphingolipids was discovered, accounting for ∼20% of the sedimentary lipidome. These sphingolipids are produced in situ by sedimentary anaerobic bacteria, which probably used sphingolipids instead of phospholipids, likely because of the deficiency of phosphate in the anoxic sediments. Our results suggest that while phytoplankton-derived lipids contribute 50–60% of the sedimentary lipidome, the importance of bacterial lipids, particularly in-situ produced sphingolipids, may have been overlooked.

## Introduction

Lipids exhibit significant structural diversity, with varied physicochemical properties reflecting their broad range of functions in organisms, including serving as membrane building blocks, energy storage molecules, signaling agents, and modulators of protein activity^1–3^. They play an important role in oceanic food webs and the carbon cycle^4–6^. In addition, intact polar lipids (IPLs)—the major molecular building blocks of cell membranes, consisting of a hydrophobic core attached to a hydrophilic polar head group—serve as biomarkers for determining the taxonomic composition of living biomass and provide insights into nutrient utilization patterns of organisms in the deep ocean^7–11^. After cell death, most IPLs lose their polar headgroups and their cores may contribute to the fossil organic matter pool^12,13^.

In marine sediments, some of these core and other lipids are more refractory in nature than DNA, proteins and sugars and can persist over long timescales, serving as biomarkers for the ancient presence of particular taxonomic groups or for reconstructing past climate conditions^4,14,15^. For example, specific isoprenoidal GDGTs (glycerol dialkyl glycerol tetraethers) can be traced as far back as the Jurassic era, and are used as biomarkers for aerobic, nitrifying archaea and for the reconstruction of past sea water temperatures^16,17^. Certain carotenoid-derived biological markers, such as isorenieratane and okenane, trace the presence of green and purple sulfur bacteria from thousands of years ago to as early as the Proterozoic period^18,19^. Additionally, C_26_ to C_30_ fossil steroids are diagnostic for Eukaryotes such as diatoms and dinoflagellates in marine environments^20–22^. On top of the influx of lipids produced in the water column, living prokaryotes in the sediments process sedimentary organic matter and possess lipid membranes that ultimately also contribute to the sedimentary lipid pool, resulting in a highly complex mixture of diverse sources.

Earlier studies of sedimentary lipids have mainly relied on analysis by gas chromatography-mass spectrometry (GC-MS), which typically limits detection to relatively small (<800 Da) and apolar lipids^1,23^. More recent applications of high-pressure liquid chromatography coupled with high-resolution mass spectrometry (HPLC-HRMS) technology have allowed for targeted analysis of lipid classes falling outside the analytical window of GC-MS, such as GDGTs, pigments, and selected intact polar lipids^23–28^. In turn, the field has gone on to develop more untargeted approaches to examine marine lipidomes, leading to the discoveries of novel lipids, as well as insights into the transport and distribution of lipids from surface waters to the deep ocean^29–35^. However, a gap remains in understanding the link between lipids that sink through the deep water column and the (sub)-seafloor sedimentary lipidome, where lipids can be stored over geological timescales. Indeed, to date, our understanding of the sedimentary lipidome remains limited to the surface sediment (i.e. < 5cm) and to a restricted range of compounds, hindering our ability to fully comprehend the contributions of all microorganisms to the lipid pool and the full lipid diversity in sediments, especially in subseafloor habitats. The advent of high-resolution untargeted lipidomic methodologies^34,36,37^, offers an opportunity to address this disparity. Here, we applied state-of-the-art untargeted lipidomic analysis of the upper 2 m of sediments from the Black Sea, resulting in a comprehensive view of the sedimentary lipidome. We investigate in unprecedented detail the composition of the sedimentary lipidome, and compare its composition with lipidomic data of suspended particulate matter (SPM) in the water column.

## Results and Discussion

### Identification and classification of the predominant lipids

We employed a recently developed analytical and computational workflow^34^ to analyze the lipidome of a sedimentary record (2 meter length, spanning sedimentation throughout the past 18 kyrs) from the Black Sea and compared this with the lipidome of the SPM in the water column (50—2000 meters below sea level, mbsl), which we studied recently ^38^. A combined molecular network of both lipidomes was constructed with the aim of assessing the diversity and abundance of lipids in both the sediments and the water column. This network was built from the clustering of ion components obtained from UHPLC-MS^*n*^ analyses of total lipid extracts, based on their structural similarities^39,40^. In total, 13,297 ion components were detected and clustered into >100 subnetworks, with 4,194 ion components classified as lipids appearing in dozens of subnetworks (Fig. 1A and Supplementary Figs. S1-2). Most of the lipids clustered in the subnetworks were chemical analogs, sharing either an identical polar headgroup (e.g., IPLs like phospholipids) or a similar core structure (e.g., alkenones, pigments), and differing by simple structural modifications such as chain length, degree of unsaturation, or hydroxylation^34^.

**Fig. 1.**
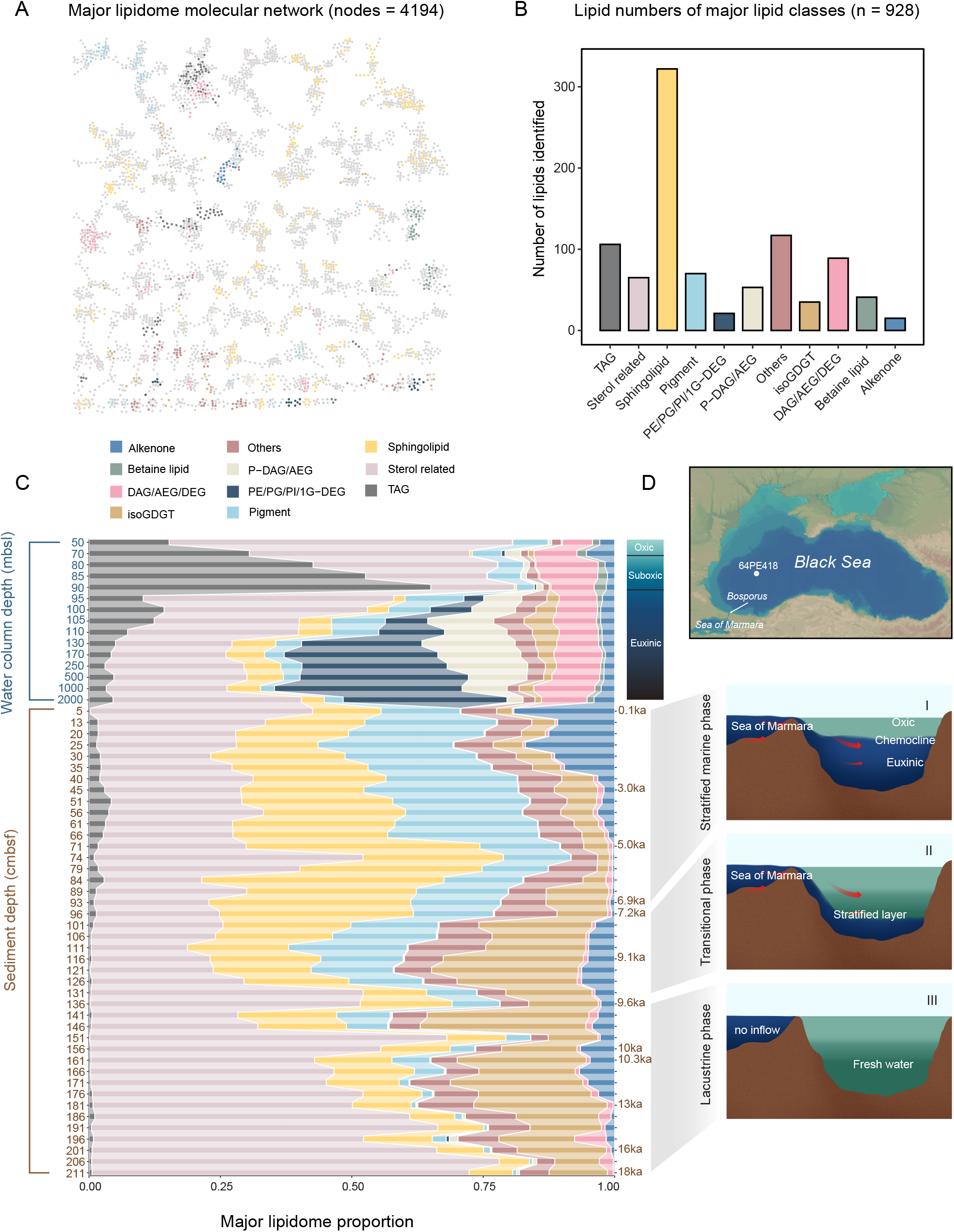
Diversity and composition of lipidomes across the water column (50–2000 m below sea level, mbsl) and sediments (5–211 cm below seafloor, cmbsf) of the Black Sea. (A) Molecular network of the lipidome, showing lipids classified based on their structural similarities. Colored nodes represent 928 major lipids that were putatively identified from the major classes shown in (B). (B) The number of individual lipids identified in the ten most abundant lipid classes and others. (C) Proportion of major lipid classes from surface water to the bottom sediment. Lipid proportions were calibrated using the response factor of external standards (Materials and Methods). Major lipid classes with a relative abundance of less than 1% in multiple samples were classified as ‘other’. (D) Map of the Black Sea basin, showing the location of core 64PE418, the stratified water column (from oxic to euxinic), and the three distinct sediment phases, with the age model based on color and elemental variations. Lipid abbreviations: 1G-DEGs (monoglycosyldietherglycerols), DAGs (diacylglycerols), DEGs (dietherglycerols), AEGs (acyletherglycerols), TAGs (triacylglycerol), PC (phosphatidylcholine), PE (phosphatidylethanolamine), PG (phosphatidylglycerol), PI (phosphatidylinositol), isoGDGT (isoprenoid glycerol dialkyl glycerol tetraethers), others include less abundant lipid classes such as SQDG (Sulfoquinovosyl diacylglycerol), BHP/BHP ester (bacteriohopanepolyols and their esters), MGDG (monoglycosyldiacylglycerol), OL (ornithine lipid), brGDGT (branched glycerol dialkyl glycerol tetraethers), diols, long chain fatty acids, quinones and etc (Supplementary Fig. S3).

A total of 928 of the most abundant lipid species were putatively identified based on accurate mass measurements of their molecular ions, MS^2^ fragmentation patterns, or comparison with data from previous studies (Fig. 1B). Overall, these lipids spanned 10 major classes: triacylglycerols (TAGs), sterols and sterol esters, sphingolipids, pigments (chlorophylls, carotenoids, sterol chlorin esters, and their alteration products), phospholipids [phosphatidylcholine (PC), phosphatidylethanolamine (PE), phosphatidylglycerol (PG), phosphatidylinositol (PI)] with varying core components [i.e., diacylglycerols (DAGs), dietherglycerols (DEGs), and “mixed” acyl/ether glycerols (AEGs, containing one ester-bound fatty acid and one ether-bond alkyl chain)], betaine lipids, glycolipids [monoglycosyldiethers (1G-DEGs)], isoprenoid glycerol dialkyl glycerol tetraethers (isoGDGTs), core lipids (DAGs/AEGs/DEGs) and alkenones. Additionally, several minor lipid classes and unknown lipid compounds were detected (Supplementary Figs. S1 and S3). The 928 identified lipids represented 75 ± 5% (across all the samples) of the total abundance from the pool of 4,194 lipids (based on peak intensity, Supplementary Fig. S2). Given the substantial differences in ionization response between lipid classes, particularly those IPLs with varying polar headgroups, sterols and pigments^37,38^, we quantified the absolute abundance of the 928 major annotated lipids using the response factors of authentic standards representative for the different lipid classes (Supplementary Tables S1-2).

### Reassessment of the lipidome composition of the Black Sea water column

Compared with our previous study of the Black Sea water column^38^, the new coupled water column-sediment molecular network contained 2 times more ion components. This increase is primarily due to methodological developments (absolute quantification and improvements in library identification), which led to the inclusion of sterols, pigments and their alteration products. Additionally, certain microbial lipids were abundant in the sediment and not in the water column, so had not been included in our previous study of the water column.

The relative abundance of sterols and TAGs was highest in the oxic and suboxic zones (50– 90 mbsl) of the water column (Fig. 1C). Sterols made up 65% of the total lipids at 50 mbsl but decreased to 16% at 90 mbsl. They likely originate primarily from phytoplankton, where they play an important role in membrane fluidity and stability^21,41^. The relative abundance of TAGs amounted to 15–65%, with the maximum at 90 mbsl. TAGs are efficient energy storage lipids produced by phytoplankton^42^. These findings are in good agreement with previous studies that investigated lipids in SPM or sinking particles from the open ocean^42,43^. Long-chain alkenones (C_36_ to C_40_), specific constituents of haptophyte microalgae and important biomarkers for paleoclimate reconstructions^44^, accounted for 3% of the total lipids in the oxic and suboxic zones.

In the euxinic part of the water column, phospholipids, primarily PE/PG/PI-DEGs, along with PE-DAGs and PE-AEGs, are dominant, comprising 11–45% of the total lipids, indicating significant anaerobic bacterial activity in deep-water environments^29,38^. Phytoplanktonic lipids, particularly sterols, remained one of the most abundant lipid groups, constituting 30 ± 8% (mean ± SD) of the total lipids. In comparison, TAGs and alkenones accounted for 7 ± 4% and 2 ± 1%, respectively. Sphingolipids, consisting of a sphingosine backbone linked to a fatty acid through an amide bond (Fig. 2A), were nearly absent in the upper oxic and suboxic waters (< 0.3%, 50–90 mbsl), but accounted for 2–8% of the total lipids in deeper waters (Figs. 1C and 2B), making them the fourth most abundant lipid class. Although this lipid class has been linked to algae and associated viruses^45,46^, our recent work, integrating metagenomic and lipidomic analysis, indicated an anaerobic bacterial origin of sphingolipids in the euxinic zone^38^.

**Fig. 2.**
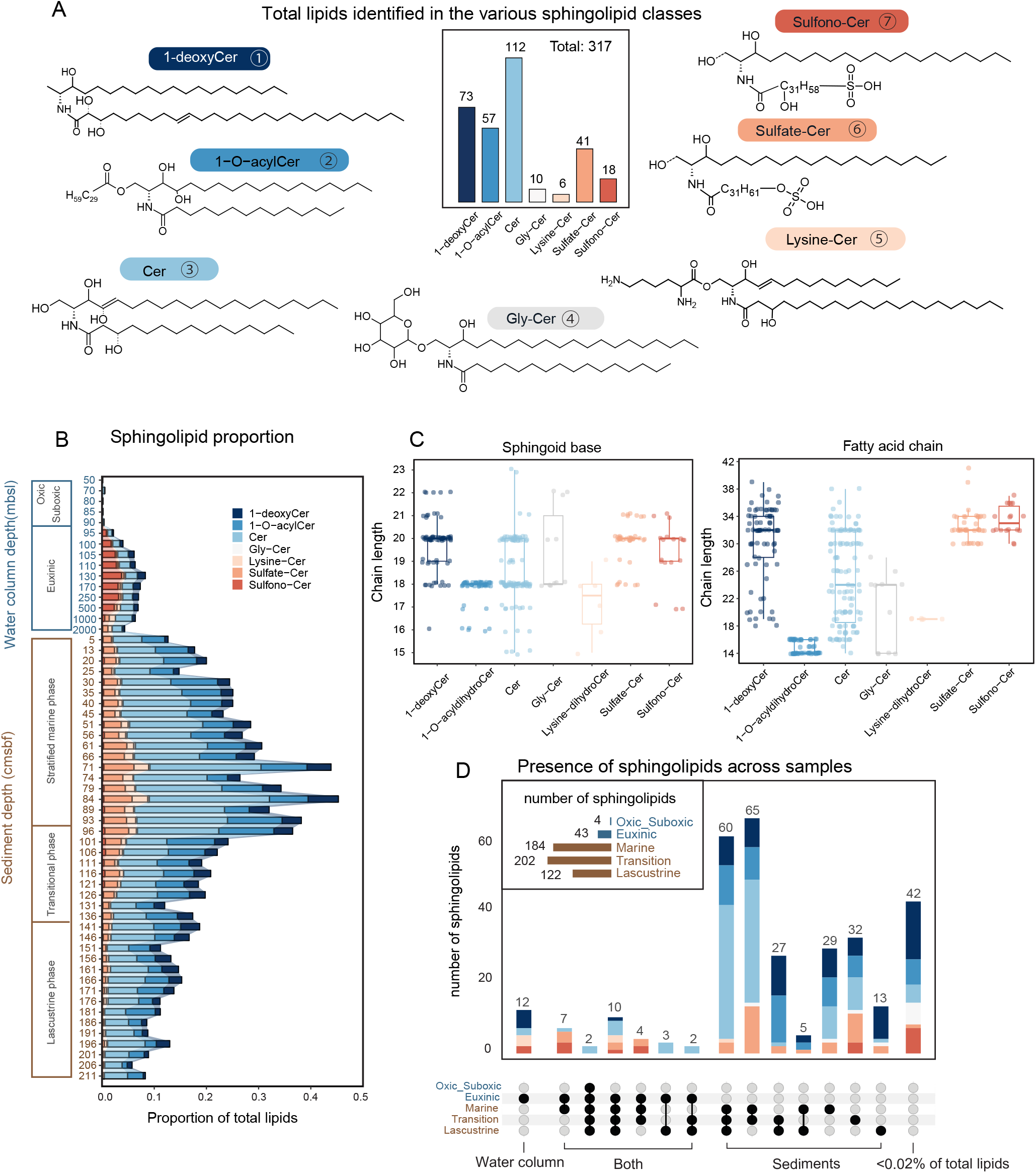
Sphingolipids in the Black Sea. (A) General structures of the 7 major sphingolipid subclasses and total of lipids identified in the various sphingolipid classes. (B) Proportion of sphingolipid subclasses to the quantified lipidome. (C) Range of carbon atom chain lengths in the sphingoid base and fatty acid for various sphingolipid subclasses. (D) Number of individual sphingolipids present in the water column and sediments, illustrated potential sphingolipid sources, whether planktonic or in-situ. A threshold of 0.02% relative abundance of total lipids was used to determine the presence of a lipid in a specific sample, shown in dark dots. Lipids with less than 0.02% total lipid abundance were shown in grey dots in that sample, connected dots denote lipid species shared across samples. Sphingolipid abbreviations: ceramides (Cer), 1-O-acyl ceramides (1-O-acyl Cer), glycosylated ceramides (Gly-Cer). For the Sulfono-Cer and Sulfate-Cer classes, the exact position of the sulfono and sulfate groups on the fatty acid chain cannot be determined from mass spectra but are identified to be located exclusively on the fatty acid chain. Additionally, dashed lines linked to OH groups indicate that these subclasses may contain OH groups.

### Major lipids in the sedimentary pool and their potential sources

The studied sediment of the Black Sea can be divided into distinct sections, representing three different phases of ancient sediment deposition (Fig. 1D): the oxygenated lacustrine phase [136– 211 cm below sea floor (cmbsf), 9.6–18 ka], the transitional phase marked by saltwater inflow from the Sea of Marmara (96–136 cmbsf, 7.2–9.6 ka), and the modern day stratified marine phase (5–96 cmbsf, 0.1–7.2 ka) with a partially euxinic water column^47^.

Our untargeted lipidomic analyses revealed that sterols and their esters accounted for 38 ± 17% (mean ± SD) of the total lipid pool and were especially abundant in the lacustrine section (Fig. 1C). They likely originate from freshwater algae and aquatic plants^48^ in the surface waters of the lake, and are thus necessarily fossilized lipids that were produced at the time of deposition. The summed concentrations ranged from 10 to 2,500 µg g^−1^ TOC in subseafloor sediments (5–211 cmbsf), consistent with previous studies using targeted methods^48,49^. Pigments and their alteration products, including chlorophylls, pheophytins, sterol chlorin esters, carotenoids and carotenoids degradation products, all predominantly attributed to phytoplankton sources, also occurred in high relative abundance (Fig. 1C). Unlike sterols, phytoplanktonic pigments and their alteration products were comparatively enriched in the marine and transitional sediment sections, comprising 13 to 29% of total lipids (mean concentration 82 µg g^−1^ TOC). In contrast, in the lacustrine section, these pigments and their alteration products accounted for 4 ± 3% of total lipids (mean concentration 54 µg g^−1^ TOC), perhaps due to the increased preservation under anoxic water column conditions. Long-chain alkenones accounted for 3 ± 2% (1 to 130 µg g^−1^ TOC) of the total lipid pool during the lacustrine and transitional phases. They were nearly absent in the early part of the marine phase but increased to 10–20% (mean 30 µg g^−1^ TOC) in the late part of the marine phase (5–35 cmbsf), in line with the fact that the marine section is primarily composed of carbonate derived from fossil coccolithophorid skeletons^50^. The fact that all these phytoplanktonic lipids, which unambiguously find an origin in the oxic surface waters, are still present in substantial abundances in the sediment confirms earlier findings that they are relatively recalcitrant and preserve well over geological time scales^19,51^.

In contrast to lipids derived from phytoplankton, bacterial phospholipids dominant in the SPM of anoxic deep waters, were nearly absent (<0.2% of total lipids) in the sedimentary lipid pool (Fig. 1C). This discrepancy may arise from two main factors. Firstly, phytoplankton lipids can rapidly be exported to the seafloor via zooplankton fecal pellets and phytoplankton aggregates, while bacterial phospholipids produced below the chemocline likely lack such a transport mechanism to the sediments. This has been demonstrated for GDGTs produced by anaerobic methane oxidizing archaea in deep anoxic waters which were not incorporated into sinking particles and hence not delivered to the sediment^52^. Secondly, phospholipids may be preferentially recycled and utilized over non-phosphorus phytoplanktonic lipids. Incubation experiments and theoretical models have demonstrated that phospholipids degrade rapidly and have limited preservation potential in sediments^12,13^. Recent work by Behrendt et al.^5^, using nano-lipidomics on heterotrophic bacteria isolated from marine particles, revealed specific members of bacterial interaction can accelerate phospholipids’ degradation. Furthermore, phospholipid recycling by bacteria leads eventually to the release of phosphorus into the water column, where it can again be taken up by eukaryotes and bacteria^53^.

### An abundant component of the sedimentary lipidome: sphingolipids

We previously reported 229 sphingolipid-like molecules in the euxinic waters of the Black Sea, with half of them putatively identified^38^, including unique 1-deoxysphingolipids with long-chain fatty acids and sulfur-containing groups. Here, in the integrated lipidome molecular network, we discovered nearly 1,000 sphingolipid-like molecules from over 20 molecular subnetworks in both the Black Sea sediments and water column (Fig. 1A). Among these, 317 distinct sphingolipid structures were putatively identified (Fig. 2A and Supplementary Figs. S4-7), representing more than one-third of the annotated lipid pool (Fig. 1B). These include some novel sphingolipid classes such as 1-O-acyl ceramides (1-O-acyl Cers) and multihydroxylated Cers. Moreover, sphingolipids were the second most abundant lipid class in the sediments, comprising 6–46% (mean ± SD: 21 ± 10%) of the total lipid pool (Fig. 1C). They were most abundant in the sediments of the stratified marine phase and, to a lesser extent, in those deposited during the transitional phase.

We classified the sphingolipid structures into seven subclasses: 1-deoxyCers (and their hydroxylated derivatives, 73 species), 1-O-acyl Cers [long-chain fatty acid (≥ C30) linked to Cer core structures, 57 species], Cers (including saturated and hydroxy Cers like phytoCers, 112 species), Glycosylated Cers (Gly-Cers, 10 species), Lysine-Cers (6 species), Sulfono-Cers (with sulfono groups linked to 1-deoxyCers and Cers, 41 species), and Sulfate-Cers (with sulfate groups linked to 1-deoxyCers and Cers, 18 species) (Fig. 2A). While the most abundant sphingolipids in the SPMs of the euxinic waters were Sulfono-Cers (2.3 ± 1.2%), Cers (2.0 ± 0.5%), and 1-deoxyCers (0.9 ± 0.4%), in the sediment core they were mainly comprising Sulfate-Cers (3.2 ± 1.1%), 1-O-acylCers (7.3 ± 2.0%), long-chain 1-deoxyCers (2.8 ± 1.1%), and Cers (12.6 ± 5.1%) (Fig. 2B). These sphingolipids assemblages displayed the widest range of fatty acid and sphingoid base chain lengths ever reported in the environment, with fatty acids ranging from 14 to 42 carbon atoms and sphingoid bases from 15 to 23 carbon atoms (Fig. 2C). Compared to other abundant classes, 1-O-acylCer had relatively shorter and narrower chain lengths, with sphingoid bases of 16–18 carbons and fatty acids of 14–16 carbons. However, the additional fatty acid linked to the C1-OH position of 1-O-acylCer contained 28–32 carbon atoms.

Examination of the number of lipids shared among samples (Fig. 2D) showed that 220 sphingolipid species were found exclusively in sediments of which 60 species were present in all three sedimentation phases, while 126 species were detected in either the marine or transitional phase, or both. Less than 30 species appeared in both the water column and sediments, such as specific Cer and Sulfono-Cer lipids (Fig. 2B). Hence, this analysis reveals a marked contrast in sphingolipid distribution between euxinic waters and sediments, suggesting that sedimentary sphingolipids are predominantly produced in-situ, with only a minor fraction exported from euxinic waters, if at all. This would be in good agreement with the fate of bacterial phospholipids in the water column, which lack efficient transport mechanisms for deposition into underlying sediments. Hence, we conclude that they represent membrane lipids of active bacteria thriving in the surface, and possibly, deeper sediments and, thus, reflect anaerobic in-situ production in the sediments. Sphingolipids have not been previously identified as an important component of the sedimentary lipidome and have only been revealed thanks to our untargeted comprehensive lipidomic approach.

To further confirm in-situ production of the sphingolipids in the sediment, we queried the Spt gene, which encodes a key enzyme for the biosynthesis of sphingolipids^38^, in publicly available metagenomes obtained from Black Sea sediments^54^ spanning the same geological time period as studied here. Even though these metagenomes do not have the deep sequencing depth required to characterize the entire sphingolipid-producing community (via metagenome-assembled genome reconstruction as in our recent study^38^), this screen revealed the presence of distinct Spt genes throughout the sediment (Supplementary Figs. S8 and S9). Because of the fast degradation rate of DNA compared to lipids, the identification of Spt genes in the sediment confirms our hypothesis of the potential of in-situ production most, if not all, sphingolipids.

This raises the question of what the primary sedimentary sources are for these compounds. Long-chain sphingolipids have been shown to be primarily produced in the euxinic water column of the Black Sea by anaerobic bacteria, such as *Desulfobacterota, Bacteroidota, Marinisomatota*, and *Chloroflexota*^38^, many of which play key roles in nitrogen and sulfur cycling^55^. These sphingolipids are linked to oxidative stress response, cell wall remodeling, and nitrogen metabolism. Another study based on prokaryotic small subunit ribosomal RNA genes and qPCR analyses^56^ showed that the classes *Anaerolineae* and *Caldilineae* within the phylum *Chloroflexota* were the most abundant bacteria in subsurface marine sediments of the Black Sea. Our metagenome data from the water column revealed that 218 *Chloroflexota* species, including those from these two classes, possess the Spt gene to synthesize sphingolipids^38^. This suggests that *Chloroflexota* could be a potential source of sphingolipids in the sediments. *Chloroflexota* is one of the most abundant bacterial communities in global marine sediments, particularly under anoxic conditions^57^. Therefore, sphingolipids produced by *Chloroflexota* might be an underappreciated class of lipids in sedimentary environments worldwide. Indeed, one of the Spt hits from Black Sea sediments is likely from a *Chloroflexota* genome, although we also report hits in other bacterial phyla (Supplementary Figs. S8 and S9).

The remarkable dominance of sphingolipids over phospholipids produced by sedimentary bacteria may be due to a lack of suitable P-substrates in the sediment^58^, leading to a preference for sphingolipid synthesis over phospholipid synthesis, as has been observed for other non-P containing lipids in low-P systems^59^. In addition, the greater stability of the amide bond in sphingolipids compared to the phosphate ester bond in phospholipids may also contribute, i.e. both lipid types may be produced in the surface sediment but P-lipids are preferentially utilized in the uppermost cm. Culturing studies of sedimentary potential sphingolipid-producers, e.g., *Chloroflexota* under phosphorus-controlled conditions may provide further insights into these hypotheses.

Regardless of the exact sources, our comprehensive analysis of the lipidome of the sediments of the Black Sea, shows that although phytoplankton-sourced lipids account for 50–60% of the total lipids in the sedimentary pool, the contribution of bacterial sphingolipids is significant, which has been overlooked in the past. These bacterial sphingolipids make up over 20% of the total sedimentary lipid pool, highlighting a significant role of bacterial in-situ production in these relatively shallow sediments. Expanding these findings to the open ocean will be essential for assessing the global sedimentary lipidome and the mechanisms that govern it.

### Methods Sampling

The piston core *64PE418*, containing approximately 210 cm of sediment, was collected from a depth of about 2000 meters below sea level (mbsl) in the western gyre of Black Sea (42°56’N, 30°02’E) during an April 2017 cruise aboard the R/V Pelagia. A total of 42 sediment samples were taken at 5 cm intervals along the core’s depth and freeze-dried for lipidome analysis. A previous study has reported the sediment chemistry, including total organic carbon (TOC) and total nitrogen (TN) content^47^. In summary, bulk TOC values range from 0.3% to 22.8%, while bulk TN values range from 0.05% to 1.9%. Accelerator Mass Spectrometry ^14^C dating of bulk organic matter was performed to establish a depth-based chronology for the core^47^.

Suspended particulate matter (SPM) was collected from the water column of the Black Sea at 15 depths ranging from 50 to 2000 mbsl during the PHOXY cruise (June 9–10, 2013). Sampling was conducted at station PHOX2 (42°54’N, 30°41’E), also within the western gyre of the Black Sea^60,61^. SPM was collected using McLane WTS-LV in-situ pumps (McLane Laboratories Inc., Falmouth), filtering through pre-washed 142-mm diameter glass fiber GF/F filters with a pore size of 0.7 µm (Pall Corporation, Port Washington, NY).

The lipidome of SPMs in the Black Sea water column has been previously reported^34,38^. In this study, we integrated these data with the newly generated lipidome of the sediment core for a more in-depth analysis. Although the SPM samples were collected from a different station, approximately 26 nautical miles from the sediment core site, the lipidome profile of the stratified water column SPMs remained relatively stable in the Black Sea^33^, allowing for meaningful comparison. It is important to note that, unlike particulate matter captured in sediment traps^43^, the lipidome from SPMs collected using an in-situ pump represents a snapshot of conditions at the time of sampling, rather than long-term sinking particulate matter. Nevertheless, it provides valuable insights into potential lipid distribution, transport within the water column, and deposition at the seafloor.

### Lipid extraction

A modified Bligh-Dyer procedure was used to extract lipids from the sediment samples^7,33^, with blank controls processed and analyzed in parallel. Both the sediment samples and blanks were subjected to ultrasonic extraction for 10 minutes, first with a mixture of methanol, dichloromethane, and phosphate buffer (2:1:0.8, v:v:v) for two rounds, followed by two extractions using methanol, dichloromethane, and aqueous trichloroacetic acid solution at pH 3 (2:1:0.8, v:v:v). The organic phase was separated by adding more dichloromethane and buffer to reach a final solvent ratio of 1:1:0.9 (v:v:v), then re-extracted three times with dichloromethane and dried under nitrogen gas. The dried extract was redissolved in a methanol solution (9:1, v:v) with an internal deuterated betaine lipid standard {1,2-dipalmitoyl-sn-glycero-3-O-4′-[N,N,N-trimethyl(d9)]-homoserine; DGTS-d9, Avanti Lipids}. Aliquots were then filtered through 0.45 µm pore size, 4 mm diameter regenerated cellulose syringe filters (Grace Alltech).

### Lipid analysis

Lipid extract analysis was conducted using UHPLC-HRMS^2^, based on the reversed-phase method of Wörmer et al.^62^ with modifications^33^. An Agilent 1290 Infinity I UHPLC system, coupled to a Q Exactive Orbitrap MS (Thermo Fisher Scientific, Waltham, MA), was used for separation. An Acquity BEH C18 column (Waters, 2.1 × 150 mm, 1.7 μm) maintained at 30°C facilitated separation. The eluent composition was: (A) MeOH/H2O/formic acid/14.8 M NH_3_aq [85:15:0.12:0.04 (v:v)] and (B) IPA/MeOH/formic acid/14.8 M NH_3_aq [50:50:0.12:0.04 (v:v)].

The elution program started with 95% A for 3 minutes, followed by a linear gradient to 40% A at 12 minutes, and 0% A at 50 minutes, which was maintained until 80 minutes. The flow rate was set at 0.2 mL min^−1^. Lipids were analyzed within an m/z range of 350–2000, with a resolving power of 70,000 ppm at m/z 200, followed by data-dependent MS^2^ with a resolving power of 17,500 ppm. The Q Exactive was calibrated to a mass accuracy of 1 ppm using Thermo Scientific Pierce LTQ Velos ESI Positive Ion Calibration Solution. Dynamic exclusion was applied during the analysis to temporarily exclude masses for 6 seconds, allowing for the selection of less abundant ions for MS^2^.

### Lipid data processing and molecular networking

The output data generated from the UHPLC-HRMS^2^ analysis were processed using MZmine software to extract MS^1^ and MS^2^ spectra and quantify peaks^63^. The processing workflow involved several key steps: mass peak detection, chromatogram building, deconvolution, isotope grouping, feature alignment, and gap filling as well as a manual check (https://ccms-ucsd.github.io/GNPSDocumentation).

The processed MS^2^ dataset was further analyzed using the Feature-Based Molecular Networking (FBMN) method through the Global Natural Product Social Molecular Networking (GNPS) platform to construct molecular networks of the detected components^39,40^. Molecular networking is a data analysis technique used in untargeted metabolomics based on MS/MS data, where MS/MS spectra are organized into a network, grouping molecules with similar spectral patterns to indicate structural similarities. Vector similarities were calculated by comparing pairs of spectra with at least six matching fragment ions, considering both the relative intensities of the fragment ions and the differences in precursor m/z values.

The molecular network was generated using MATLAB scripts, allowing each spectrum to connect to its top K scoring matches (typically up to 10 connections per node). Connections (edges) between spectra were retained if they were among the top K matches for both spectra and if their vector similarity score (cosine value) exceeded a user-defined threshold. In this study, a cosine threshold of 0.6 was used, where a cosine value of 1.0 represents identical spectra^64^. The resulting molecular network was then imported into Cytoscape for visualization and further analysis^65^.

Each node in the molecular network stands for an individual molecule associated with a specific MS^2^ spectrum. Due to the extraction and analytical methods, and based on annotations of the similar data in a previous study^38^, most of the ion components from the molecular network we detected were lipids, thus we used the term lipidome where the ion components are discussed. Most of the molecules that clustered together in the subnetworks were either analogs of each other with an identical headgroup or with a similar core, differing by simple transformations such as alkylation, unsaturation, and glycosylation.

### Lipid standard calibration

Calibration of lipid concentrations using external standards is crucial to account for significant differences in ionization response between lipid classes, particularly those with varying polar headgroups^37,38^. Supplementary Tables S1 and S2 detail the standards used for quantifying each lipid class. It was assumed that all lipids within the same class have similar response factors, so calibration curves were generated using a single representative lipid species for each class. For sphingolipids, including ceramides (Cer) and lysine-ceramides (Lysine-Cer), an average response factor from a ceramide mix was applied (Supplementary Tables S1 and S2). Sphingolipids associated with 1-deoxyCer were calibrated using the external standard 1-deoxyCer (d18:1/24:0), while glycosphingolipids (Gly-Cer) were calibrated using Glucosyl (β) C12 Cer. Five sterols— Epicoprostanol, Coprostanol, Stigmasterol, Cholesterol, and β-Sitosterol—were used to calibrate sterol-related lipids. The most abundant ion in these sterol standards was [M+H-H_2_O]^+^, and their peak intensities were quantified using the sum of [M+H]^+^, [M+NH_4_]^+^, [M+Na]^+^, and [M+H-H_2_O]^+^ adducts.

Chlorophyll a was used to quantify all pigment-related lipids, including chlorophyll, bacteriopheophytin, sterol chlorin esters, carotenoids and their degraded products. As reported previously^43^, chlorophyll a was detected as pheophytin a, likely due to demetallation during ionization or due to the eluent composition. Therefore, we used the peak intensity of pheophytin a to quantify the chlorophyll a standard, acknowledging that chlorophyll a was likely the dominant source of this signal. None of the other lipid standards showed signs of degradation. For lipid classes without specific standards, such as ornithine lipids (OLs), a response factor from another class of amino lipids (DGTS) was used. Unknown lipid classes were calibrated using the average response factor of all external standards.

Lipid concentrations were expressed as ng L^−1^ for SPM samples from the water column and ng g^−1^ or µg g^−1^ TOC for sediment samples. The proportion of lipid species identified was then used for further analysis.

### Lipidome annotation

Recently, we contributed the mass spectra of approximately 250 representative lipids from the major lipid classes of water column SPMs in the Black Sea to the open-access GNPS mass spectra library^34,38^. In this study, the MS^2^ spectra from the combined SPM and sediment core dataset were first searched against the GNPS library, resulting in less than 1% annotation. The most abundant lipids from each class were then annotated by comparison with data from previous studies^18,29–32,66–74^ or putatively identified based on accurate mass and MS^2^ fragmentation patterns. Based on the annotated lipids from each subnetwork, we identified the major lipid classes and used their calibrated concentration for lipidome composition analysis.

## Statistical analysis

For principal component analysis (PCA), lipid species abundance data were first transformed using the Hellinger distance method to reduce bias from zero values. The transformed data were then processed and visualized using R software (version 4.1.2). Hierarchical clustering was carried out with the “ggplot2” and “pheatmap” packages in R. An upset plot was generated using the “UpSetR” package in R^75^.

### Query of Spt in sedimentary Black Sea metagenomes

We downloaded the quality-controlled reads from MG-RAST that were reported in More et al.^54^. The reads were assembled with MEGAHIT (v1.2.9)^76^, and proteins were predicted with Prodigal in metagenomic mode (v2.6.3)^77^. Spt-like sequences were identified similarly to Ding and von Meijenfeldt et al.^38^ based on phylogenetic clustering with known Spt sequences. We used BLASTP (v2.12.0)^78^ with the same query sequences as in Ding and von Meijenfeldt et al.^38^ and the database size set to 1e8 to identify homologs. Hits with an *e*-value ≤1e-70 and query coverage ≥70 were placed in a phylogenetic tree with known Spt sequences and outgroup sequences from other alpha-oxoamine synthases (Kbl, HemA, and BioF). For known Spt sequences we used 10% of the sequences identified as clade B in Ding and von Meijenfeldt et al.^38^. The proteins were aligned with MAFFT (v7.505)^79^ using the L-INS-i algorithm, and trimmed with trimAl (v1.4.rev15)^80^ in gappyout mode. The maximum-likelihood phylogenetic tree was constructed with IQ-TREE (v2.1.2)^81^, employing 1,000 ultrafast bootstraps. Model Finder^82^ was set to nuclear models and the best-fit model (LG+R8) was chosen according to the Bayesian Information Criterion (BIC). The tree was visualized in interactive Tree Of Life (iTOL)^83^.

Contigs that contained Spt homologs were taxonomically annotated with Contig Annotation Tool (CAT) from the CAT pack software suite (v6.0.1)^84^, using a reference database based on GTDB^85^ downloaded on 20 November 2023. Prodigal in metagenomic mode (v2.6.3) and DIAMOND (v2.0.6)^86^ were used for protein prediction and alignment, respectively. CAT was run with default parameters and the DIAMOND --top parameter set to 11.

## Acknowledgments

We thank Rick Hennekam, Laura Villanueva as well as the captain and crew of the R/V Pelagia for the collection of the sediment core during Black Sea expedition-2017 (*64PE418*). We thank Marcel van der Meer and Laura Villanueva for helpful discussions. Denise Dorhout, Monique Verweij, Michel Koenen and Anchelique Mets are acknowledged for their technical support.

## Author contributions

S.D., N.B., S.S. and J.S.S.D conceived the research. S.D. analyzed the data. S.D. and N.B. identified the lipidome. A.C. performed the lipid extraction. F.A.B.v.M. performed metagenome analyses. S.D., N.B., S.S. and J.S.S.D interpreted the data. J.S.S.D. acquired funding. J.S.S.D. and S.S. supervised this project. S.D. wrote the initial manuscript with contributions from all authors.

## Data and materials availability

The mass spectra data and quantification data (.mgf and.csv) with the molecular network and detailed parameter settings can be accessed at the GNPS platform: https://gnps.ucsd.edu/ProteoSAFe/status.jsp?task=fe89ca45b3664064b394c7d64fb454b1. The source data are available at Zenodo at DOI: 10.5281/zenodo.14203114.

## Supplementary Materials

Supplementary Text

Supplementary Figs. S1 to S9

Supplementary Tables S1 to S2

