## Supplementary for "Substantial contribution of in-situ produced bacterial lipids to the sedimentary lipidome"

**The PDF file includes:**

Supplementary Text

Supplementary Figs. S1 to S9

Supplementary Tables S1 to S2

**Supplementary Text**

Identification of unassigned sphingolipids

As described in the main text, many of the sphingolipid subnetworks in our dataset contained unassigned components (Fig. 2 and Supplementary Fig. S1). Representative structural elucidations of Cer, Gly-Cer, Lysine-Cer, 1-deoxyCer, Sulfono-1-deoxyCer, and Sulfate-1-deoxyCer were presented in our previous study[34]. In this study, several new structures from these classes were identified, particularly those with additional hydroxy groups on the sphingolipid base (either the fatty acid or sphingosine). Here, we elucidate the identities of two unassigned components from newly discovered sphingolipid classes, 1-O-acyldihydroCer and Sulfate-Cer, both containing extra hydroxy groups.

The subnetwork of 1-O-acyldihydroCer contains 57 components, ranging from *m/z* 916.9065 to *m/z* 1029.0317. We focus first on a representative lipid labeled with *m/z* 948.9681 (Fig. S4 for representative $MS^2$ spectra). The $MS^2$ spectrum showed a loss of 436.4637 Da (AEC $C_{30}H_{60}O$; Δm = -0.17 mmu) and the formation of a dominant fragment ion at *m/z* 510.4877 with an AEC of $C_{32}H_{64}O_3N^+$ (Δmmu = -0.37), likely representing the C30:0 acyl chain and the sphingolipid core structure, respectively. Fragment ions at *m/z* 282.2789 with AEC of $C_{18}H_{36}ON^+$ (Δmmu = -0.24) and *m/z* 228.2319 with AEC of $C_{14}H_{30}ON^+$ (Δmmu = -0.29), confirms that the sphingolipid core is composed of C18:0 and C14:0 chains. In addition to the $C_{18}H_{36}ON^+$ fragment, $C_{18}H_{34}N^+$ and $C_{18}H_{38}O_2N^+$ fragments were also observed, suggesting that the sphingoid base comprises C18:0 chain with two hydroxy groups. Therefore, the fatty acid group of sphingolipid core is composed of C14:0 chain. We tentatively identified this component as 1-O-acyl(30:0)-dihydroCer (d18:0/14:0). 1-O-acylCers are characteristic of eukaryotic systems, particularly the outer layers of vertebrate skin, where they play a critical role in forming the lipid barrier of the stratum corneum[87]. To the best of our knowledge, this is the first time 1-O-acylCers have been found in sediments and are supposed to be produced by bacteria.

Another sphingolipid class, Sulfate-Cer, which contains largely unknown components in the molecular network, is composed of 41 components with *m/z* values between 830.6909 and 944.7944 (Fig. 2A and Supplementary Fig. S1). Fig. S6 shows the $MS^2$ spectrum of a member of this cluster with a retention time of 19.4 min and *m/z* 876.7318 (*m/z* 876.7310 in Supplementary Fig. S6), assigned an EC of $C_{50}H_{102}O_8NS^+$ (Δm = -0.81 mmu). A dominant fragment ion at *m/z* 778.7631 with an AEC of $C_{50}H_{100}O_4N^+$ is observed (Δm = -1.58 mmu), indicating that the initial neutral loss of 80 Da was $H_2SO_4$ (the sulfate moiety). A continuous loss of $H_2O$ is observed from *m/z* 778.7631 to *m/z* 724.7349, suggesting the presence of three hydroxy groups on the sphingolipid base. Fragment ions produced from the fatty acid moiety, *m/z* 724.7349 with AEC of $C_{30}H_{60}O_2N^+$ and *m/z* 431.4260 with AEC of $C_{30}H_{55}O^+$, suggest a hydroxy group located on the fatty acid chain. A loss of the C30:2 acyl chain from m/z 760.7527 results in a C20-sphingoid base (*m/z* 312.3255, AEC of $C_{20}H_{42}ON^+$). Fragments at m/z 330.3362 with an AEC of $C_{20}H_{42}O_2N^+$ indicate that this is a Cer-sphingoid base with two hydroxy groups located at the C1 and C3 positions. The sulfate moiety is likely located on the fatty acid base rather than the C1-OH sphingoid base as observed. Therefore, we tentatively identify this component as Sulfate-dihydroCer (d20:0/OH-30:0).

For all discussed components above, the structural identification is tentative. Further structural elucidation methods (e.g., nuclear magnetic resonance [NMR] spectroscopy) would be required to confirm the proposed structures and elucidate the position of the various functional groups.

**Fig. S1.**

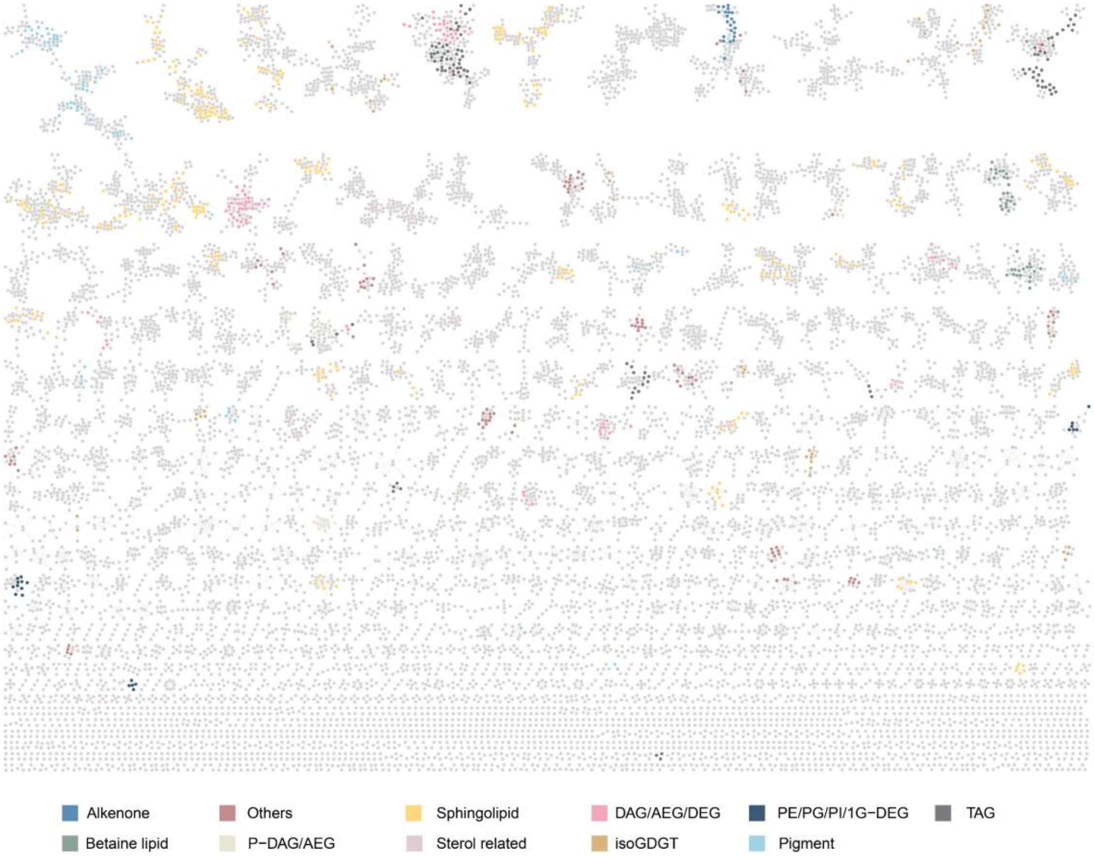


Molecular network of 13,297 ion components detected from the Black Sea water column and sediment, of which 4,194 are lipid species clustered into dozens of subnetworks (Fig. 1A). Colored nodes represent 928 major lipids putatively identified from the major classes. Lipid abbreviations: 1G-DEGs (monoglycosyldietherglycerols), DAGs (diacylglycerols), DEGs (dietherglycerols), AEGs (acyletherglycerols), TAGs (triacylglycerols), PC (phosphatidylcholine), PE (phosphatidylethanolamine), PG (phosphatidylglycerol), PI (phosphatidylinositol), isoGDGT (isoprenoid glycerol dialkyl glycerol tetraethers). Other, less abundant lipid classes include SQDG (sulfoquinovosyl diacylglycerol), BHP/BHP ester (bacteriohopanepolyols and their esters), MGDG (monoglycosyldiacylglycerol), OL (ornithine lipid), brGDGT (branched glycerol dialkyl glycerol tetraethers), diols, long-chain fatty acids, quinones, etc.

**Fig. S2.**

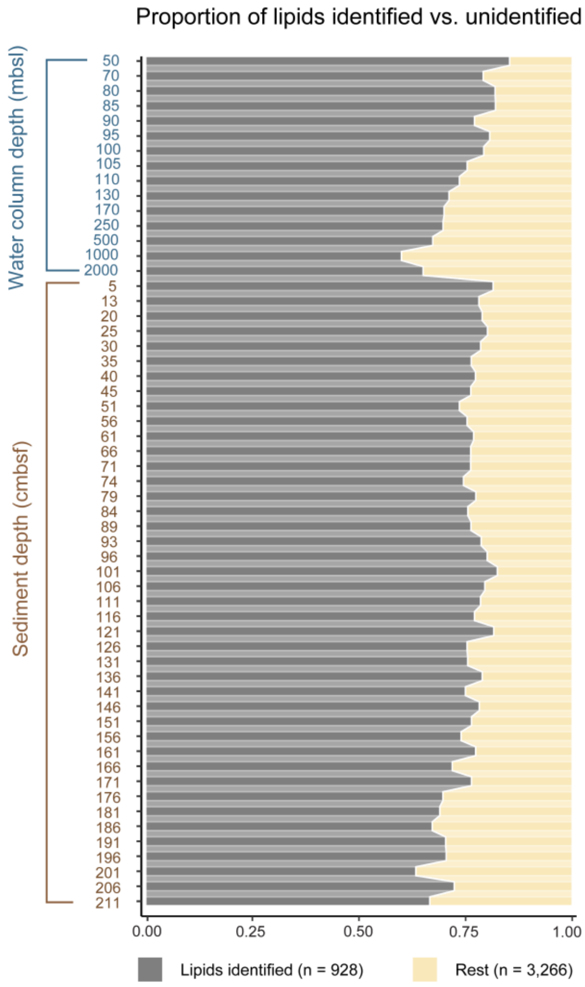


Proportion of identified lipids (n = 928) versus unidentified lipids (n = 3,266) based on peak intensity from the lipid-containing molecular network (Fig. 1A). The 928 identified lipids accounted for 75 ± 5% of the total lipids (n = 4,194, Fig. 1A) in all samples and were therefore used to represent the major lipid pool for subsequent analysis.

**Fig. S3**

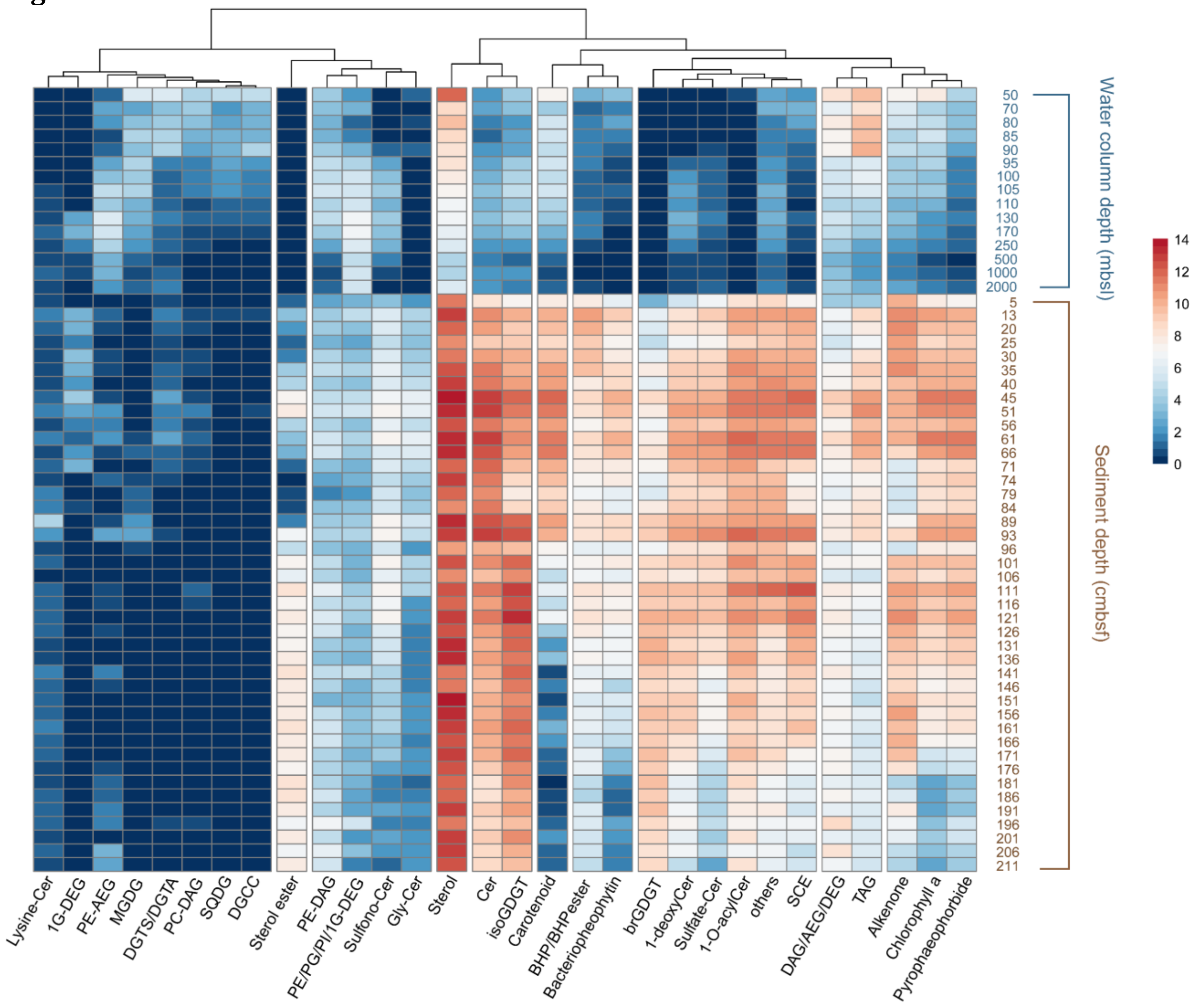


Hierarchical clustering heatmap illustrating the distribution of 29 lipid classes across the water column (50–2,000 m below sea level, mbsl) and sediments (5–211 cm below seafloor, cmbsf). The color bar on the right represents the Z-score normalization scale of lipid class abundance, calculated using the $\log_2$(1 + lipid class abundance) transformation. This heatmap includes 29 lipid classes, providing a more detailed distribution than Fig. 1B-C, which showed only the 10 most abundant classes. The extended set of lipid classes offers an overview of their distribution in the Black Sea and highlights their correlations across the water column and sediments. Lipid abbreviations: 1G-DEGs (monoglycosyldietherglycerols), DAGs (diacylglycerols), DEGs (dietherglycerols), AEGs (acyletherglycerols), TAGs (triacylglycerol), DGTS [diacylglycerylhydroxymethyltrimethyl-(N,N,N)-homoserine], DGCC (diacylglycerylcarboxyhydroxymethylcholine), DGTA (diacylglyceryl-hydroxymethyl-tri-methyl-b-alanine), MGDG (monoglycosyldiacylglycerol), PC (phosphatidylcholine), PE (phosphatidylethanolamine), PG (phosphatidylglycerol), PI (phosphatidylinositol), isoGDGT (isoprenoid glycerol dialkyl glycerol tetraethers), brGDGT (branched glycerol dialkyl glycerol tetraethers), Cer (ceramides), Gly-Cer (glycosylated ceramides), SCE (steryl chlorin esters), SQDG (Sulfoquinovosyl diacylglycerol), BHP/BHP ester (bacteriohopanepolyols and their esters), others include less abundant lipid classes such as OL (ornithine lipid), diols, long chain fatty acids, quinones and etc.

**Fig. S4.**

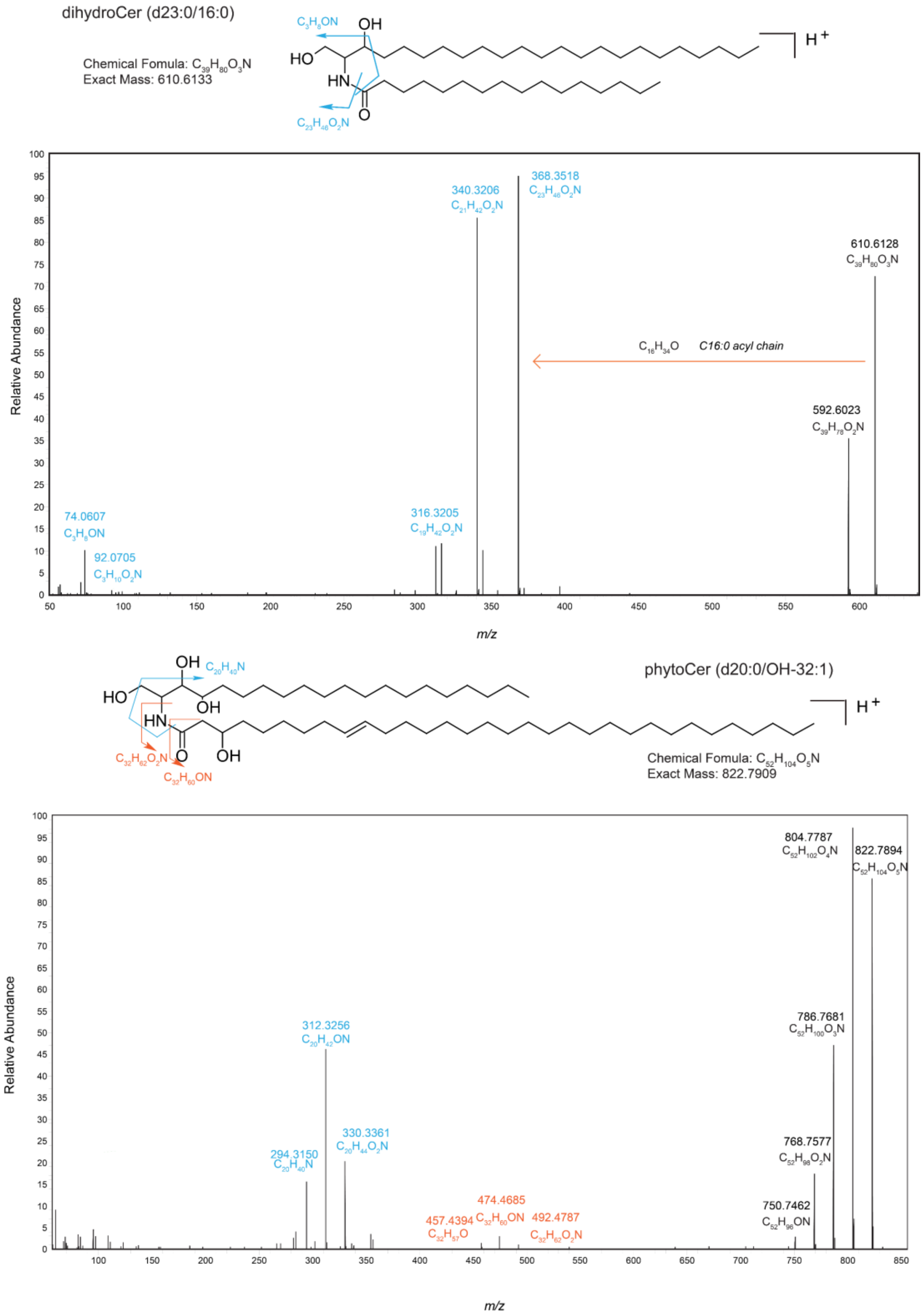


The MS[2] mass spectra and identification of dihydroCer (d23:0/16:0) and phytoCer (d20:0/OH-32:1), representative examples of structures from the Cer class.

**Fig. S5.**

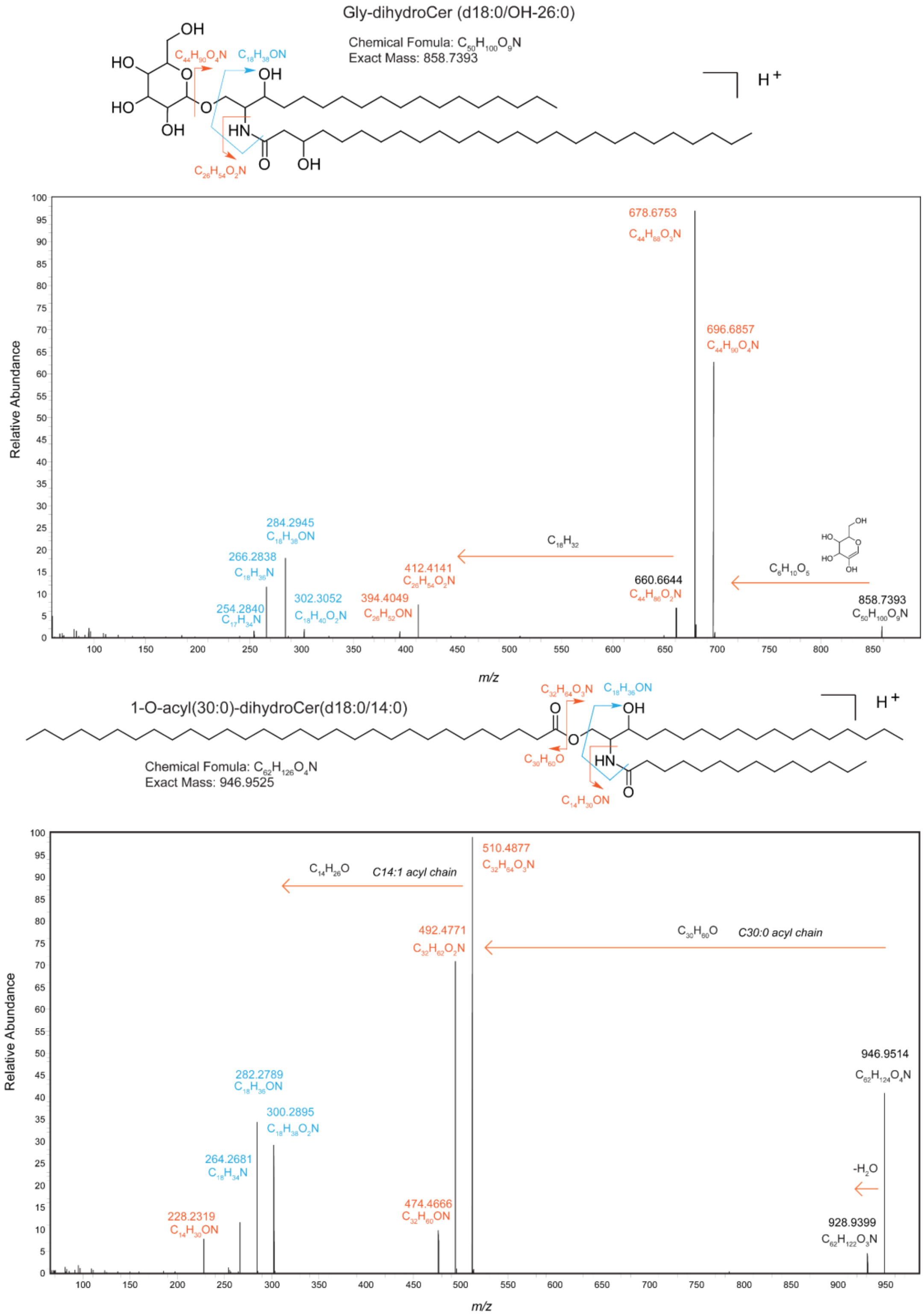


The MS$^2$ mass spectra and identification of Gly-dihydroCer (d18:0/OH-26:0) and 1-O-acyl (30:0)-dihydroCer (d18:0/14:0), representative examples of structures from the Gly-Cer and 1-O-acylCer classes, respectively.

**Fig. S6.**

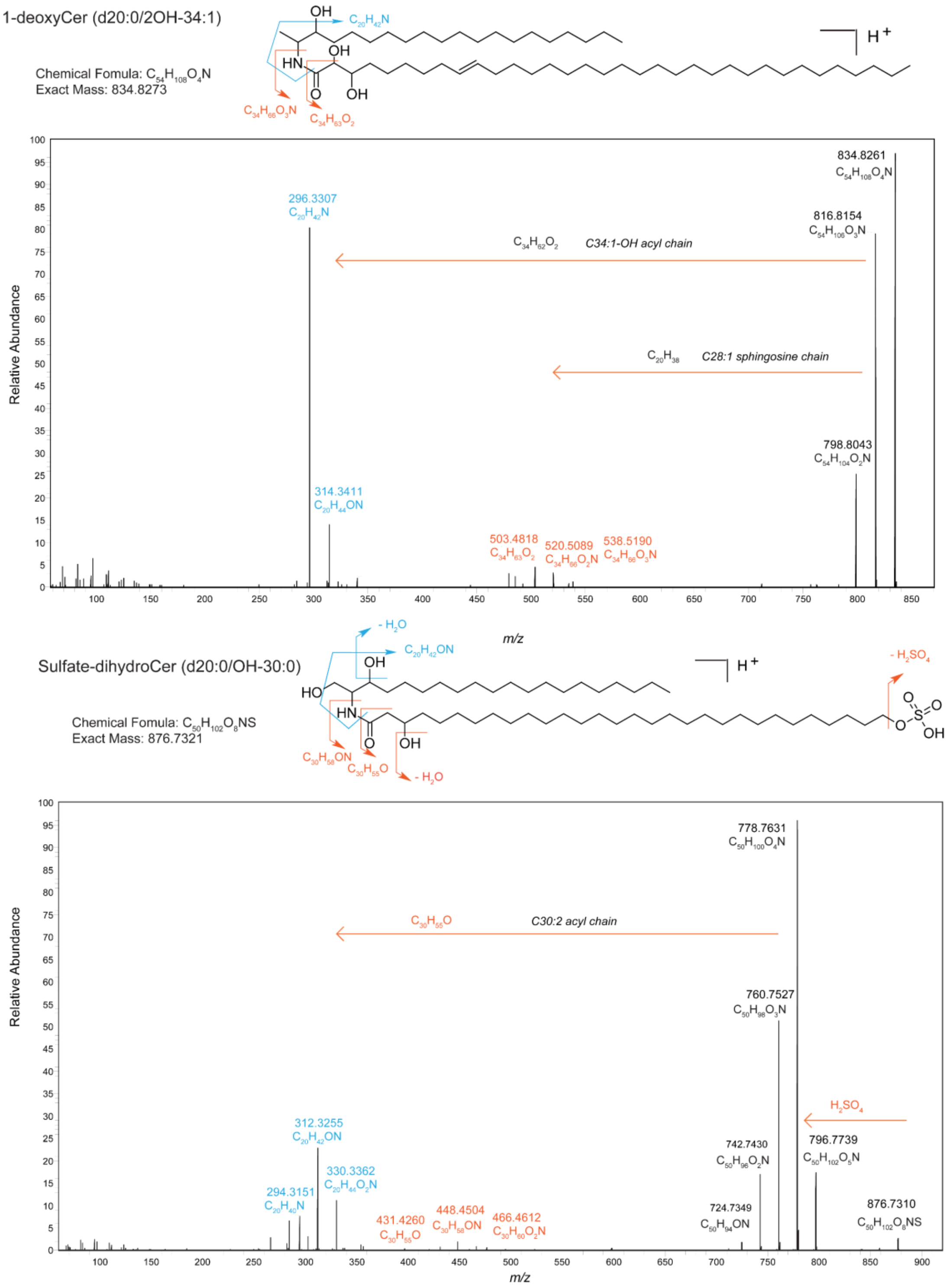


The $MS^2$ mass spectra and identification of 1-deoxyCer (d20:0/2OH-34:1) and Sulfate-dihydroCer (d20:0/OH-30:0), representative examples of structures from the 1-deoxyCer and Sulfate-Cer classes, respectively.

**Fig. S7.**

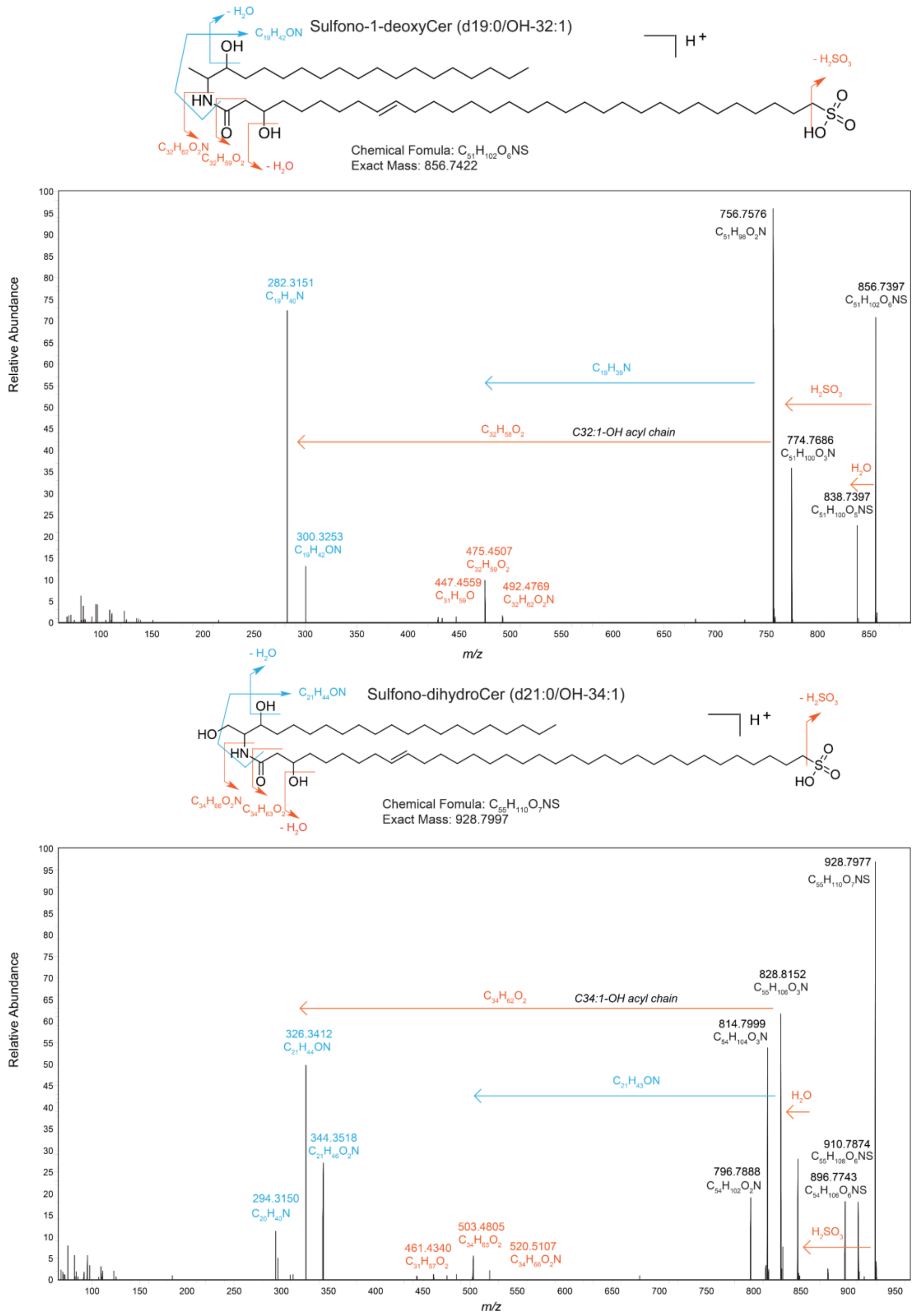


The MS[2] mass spectra and identification of Sulfono-1-deoxyCer (d19:0/OH-32:1) and Sulfono-dihydroCer (d21:0/OH-34:1), representative examples of structures from the Sulfono-Cer class.

**Fig. S8.**

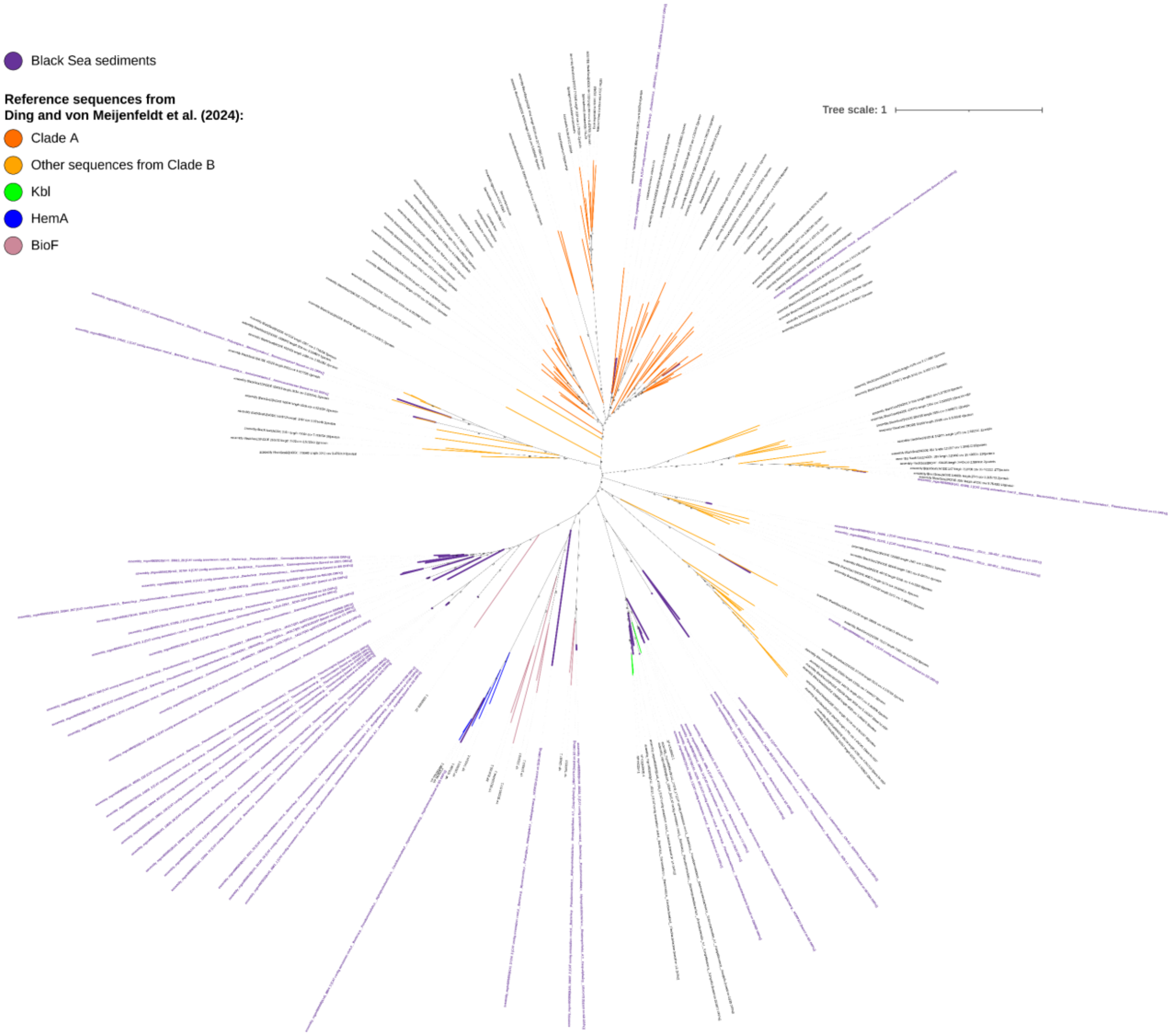


Identification of Spt homologs in Black Sea sediment. Bold purple sequences are hits from the metagenomes of Black Sea sediments reported in More et al.[54], other sequences are a selection of backbone sequences reported in Ding and von Meijenfeldt et al.[38]. All Kbl, HemA, and BioF sequences that were included in Ding and von Meijenfeldt et al.[38] are also included here. Hits from the Black Sea sediment that cluster with these sequences are likely alpha-oxoamine synthases with an enzymatic function that differs from that of Spt and not involved in sphingolipid biosynthesis. Dark orange and light orange sequences are a selection of Spt-like sequences identified in the water column of the Black Sea. 10% of hits from Fig. 3b in Ding and von Meijenfeldt et al.[38] were included. The taxonomic annotation based on CAT is present in the name of the Black Sea sediment hits. Ultrafast bootstrap support values are indicated on the branches of the tree.

**Fig. S9.**

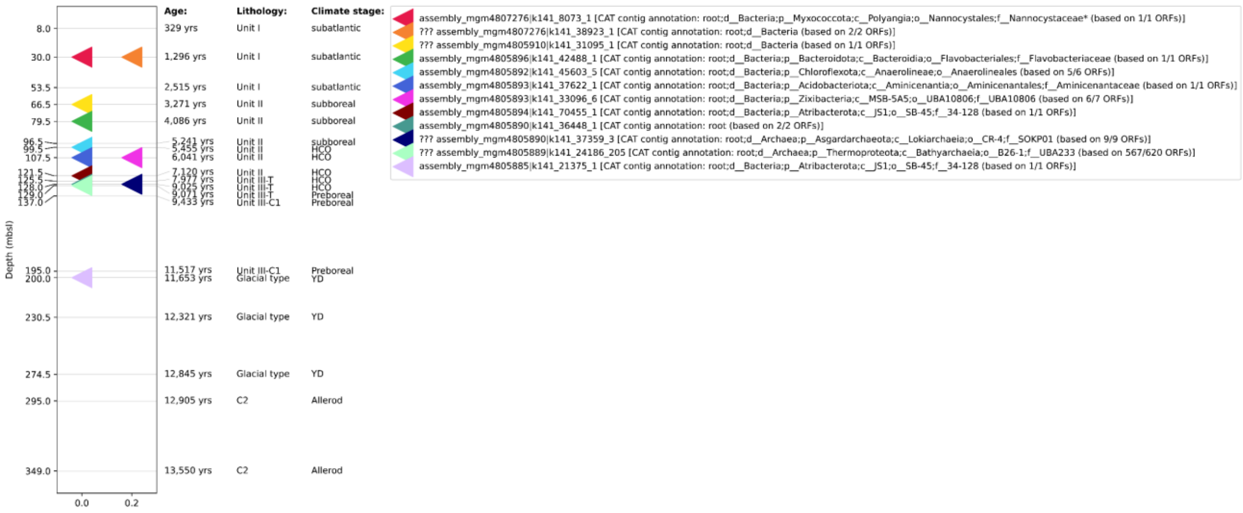


Distribution of Spt-like sequence hits in metagenomes of Black Sea sediments. Hits present in fig. S8 that are clustering with backbone Spt sequences and not with alpha-oxoamine synthases are plotted. The four hits that start with ‘???’ are clustering in between Kbl hits and the Spt-like sequences (see Fig. S8), and their phylogenetic placement is thus inconclusive to assign likely enzymatic function. Age, lithology, and climate stage of the sediment core are based on the interpretation of More et al.[54].

**Table S1.**

Lipid standards information used for quantification of major lipid classes, including their full name, molecular formula, and exact mass. All standards, except archaeol and crenarchaeol, were ordered from Avanti Polar Lipids. Archaeol and crenarchaeol were isolated from archaeal cultures in our lab.

| Standard | Full name | Molecular Formula | Exact Mass |
|---|---|---|---|
| DGTS (32:0) | 1,2-dipalmitoyl-sn-glycero-3-O-4'-(N,N,N-trimethyl)-homoserine | C42H81NO7 | 711.6013 |
| C16 ceramide (d18:1/16:0) | N-palmitoyl-D-erythro-sphingosine | C34H67NO3 | 537.5121 |
| C18 ceramide (d18:1/18:0) | N-stearoyl-D-erythro-sphingosine | C36H71NO3 | 565.5434 |
| C24 ceramide (d18:1/24:0) | N-lignoceroyl-D-erythro-sphingosine | C42H83NO3 | 649.6373 |
| C24:1 ceramide (d18:1/24:1 (15Z)) | N-nervonoyl-D-erythro-sphingosine | C42H81NO3 | 647.6217 |
| Glucosyl (β) C12 Ceramide | N-(dodecanoyl)-1-ß-glucosyl-sphing-4-ene | C36H69NO8 | 643.5023 |
| 1-deoxyceramide (d18:1/24:0) | N-[(2S,3R,4E)-3-Hydroxy-4-octadecen-2-yl]tetracosanamide | C42H83NO2 | 633.6424 |
| Hydrogenated MGDG | monogalactosyldiacylglycerol (plant, hydrogenated) | C43H82O10 | 758.5908 |
| Hydrogenated DGDG | Digalactosyldiacylglycerol (plant, hydrogenated) | C51H96O15 | 948.6749 |
| 15:0-18:1(d7) PC | 1-pentadecanoyl-2-oleoyl(d7)-sn-glycero-3-phosphocholine | C41H74D7NO8P | 752.6055 |
| 15:0-18:1(d7) PE | 1-pentadecanoyl-2-oleoyl(d7)-sn-glycero-3-phosphoethanolamine | C38H67D7NO8P | 710.5586 |
| 15:0-18:1(d7) PG | 1-pentadecanoyl-2-oleoyl(d7)-sn-glycero-3-[phospho-rac-(1'-glycerol)] (sodium salt) | C39H68D7O10P | 763.5351 |
| 15:0-18:1(d7) PI | 1-pentadecanoyl-2-oleoyl(d7)-sn-glycero-3-phosphoinositol (ammonium salt) | C42H72D7O13P | 846.5958 |
| 18:1(d7) LPC | 1-oleoyl(d7)-2-hydroxy-sn-glycero-3-phosphocholine | C26H45D7NO7P | 528.3915 |
| 18:1(d7) LPE | 1-oleoyl(d7)-2-hydroxy-sn-glycero-3-phosphoethanolamine | C23H39D7NO7P | 486.3446 |
| 18:1(d7) MG | 1-oleoyl(d7)-rac-glycerol | C21H33D7O4 | 363.3361 |
| 15:0-18:1(d7) DG | 1-pentadecanoyl-2-oleyol(d7)-sn-glycerol | C36H61D7O5 | 587.5501 |
| 15:0-18:1(d7)-15:0 TG | 1,3-dipentadecanoyl-2-oleyol(d7)-glycerol | C51H89D7O6 | 811.7641 |
| 18:1(d9) SM | N-oleoyl(d9)-D-erythro-sphingosylphosphorylcholine | C41H72D9N2O6P | 737.6392 |
| Cholesterol (d7) | cholesterol-d7 | C27H39OD7 | 393.3983 |
| 18:1 Chol ester | cholesteryl oleate | C45H78O2 | 650.5996 |
| C18:C18 wax ester | Oleic acid oleyl ester | C36H72O2 | 536.5527 |
| C32:6 FA | 14Z,17Z,20Z,23Z,26Z,29Z-dotriacontahexaenoic acid | C32H52O2 | 468.3962 |
| Epicoprostanol | 5β-Cholestan-3α-ol | C27H48O | 388.3699 |
| Coprostanol | 5β-Cholestan-3β-ol | C27H48O | 388.3699 |
| Stigmasterol | 5,22-Stigmastadien-3β-ol | C29H48O | 412.3699 |
| Cholesterol | 5-Cholesten-3β-ol | C27H46O | 386.3543 |
| β-Sitosterol | 5-Stigmasten-3β-ol | C29H50O | 414.3856 |
| chl-a | chlorophyll-a | C55H72MgN4O5 | 892.5347 |
| archaeol | archaeol | C43H88O3 | 652.6728 |
| crenarchaeol | crenarchaeol | C86H162O6 | 1291.2366 |

**Table S2.**

Lipid standards used in curves for quantification of major lipid classes. Response factors for each class of lipids were approximated by external calibration with standards. The slope of the linear curve (Peak Area/ngOC) as well as the ratio of that slope to the DGTS-d9 reference standard (Response factor relative to DGTS-d9 curve) is also reported.

| Standard | Used to quantify | Peak area/ngOC | Response factor |
|---|---|---|---|
| DGTS-d9 | | 1.58E+09 | 1.0000 |
| C16 ceramide (d18:1/16:0) | Treat as ceramide mixers, use average response factor to quantify ceramide related lipids | 1.88E+08 | 0.1184 |
| C18 ceramide (d18:1/18:0) | | 2.01E+08 | 0.1268 |
| C24 ceramide (d18:1/24:0) | | 1.64E+08 | 0.1036 |
| C24:1 ceramide (d18:1/24:1 (15Z)) | | 1.85E+08 | 0.1165 |
| Ceramide mixers | Ceramide related sphingolipids | | 0.1163 |
| Glucosyl (β) C12 Ceramide | Glycosylceramides | 1.40E+08 | 0.0883 |
| 1-deoxyceramide (d18:1/24:0) | 1-deoxyceramide related sphingolipids | 2.25E+08 | 0.1420 |
| Hydrogenated MGDG | Monogalactosyldiacylglycerol (MGDG) | 9.54E+07 | 0.0602 |
| Hydrogenated DGDG | Digalactosyldiacylglycerol (DGDG) | 1.66E+07 | 0.0105 |
| DGTS (32:0) | DGTS (diacylglyceryl trimethylhomoserines)<br>DGCC (diacylglyceryl-3-Ocarboxyhydroxymethylcholine)<br>DGTA (diacylglyceryl hydroxymethyl-trimethyl-β-alanine) | 9.43E+08 | 0.5954 |
| archaeol | Archaeol related lipids | 3.64E+07 | 0.0260 |
| crenachaeol | GDGT (Glycerol dialkyl glycerol tetraether lipids) | 1.16E+08 | 0.0830 |
| 15:0-18:1(d7) PC | Phosphatidylcholine (PC) | 6.77E+08 | 0.4272 |
| 15:0-18:1(d7) PE | Phosphatidylethanolamine (PE) | 9.67E+07 | 0.0610 |
| 15:0-18:1(d7) PG | Phosphatidylglycerol (PG) | 2.63E+07 | 0.0166 |
| 15:0-18:1(d7) PI | Phosphatidylinositol (PI) | 1.97E+07 | 0.0125 |
| 18:1(d7) LPC | LysoPC | 7.38E+08 | 0.4661 |
| 18:1(d7) LPE | LysoPE | 4.15E+07 | 0.0262 |
| 18:1(d7) MG | Monoglycerols (MGs) | 5.56E+07 | 0.0351 |
| 15:0-18:1(d7) DAG | Diacylglycerols (DAGs)<br>Acyletherglycerols (AEGs)<br>Dietherglycerols (DEGs) | 2.22E+08 | 0.1400 |
| 15:0-18:1(d7)-15:0 TAG | Triacylglycerols (TAGs) | 4.22E+08 | 0.2661 |
| C18:C18 wax ester | diols, triols and wax esters | 2.34E+08 | 0.3652 |
| C32:6 FA | long chain fatty acids and alkenones | 1.30E+08 | 0.0821 |
| Cholesterol (d7) | Treat as sterol mixers, use average response factor to quantify sterol related lipids | 1.10E+06 | 0.0010 |
| Epicoprostanol | | 2.54E+06 | 0.0016 |
| Coprostanol | | 3.87E+05 | 0.0002 |
| Stigmasterol | | 2.01E+06 | 0.0013 |
| Cholesterol | | 2.75E+06 | 0.0017 |
| β-Sitosterol | | 3.80E+06 | 0.0024 |
| sterol mixers | Sterols related lipids | | 0.0014 |
| chl-a | pigments | 1.47E+08 | 0.0930 |
| 18:1 Chl ester | sterol esters | 1.18E+08 | 0.0742 |
| average standard value | other lipids | | 0.1449 |

For sphingolipids, including Cer and Lysine-Cer, an average response factor from a ceramide mix was applied. Sphingolipids associated with 1-deoxyCer were calibrated using the external standard 1-deoxyCer (d18:1/24:0), while Gly-Cer were calibrated using Glucosyl (β) C12 Cer. Five sterols—Epicoprostanol, Coprostanol, Stigmasterol, Cholesterol, and β-Sitosterol—were used to calibrate sterol-related lipids. The most abundant ion in these sterol standards was $[M+H-H_2O]^+$, and their peak intensities were quantified using the sum of $[M+H]^+$, $[M+NH_4]^+$, $[M+Na]^+$, and $[M+H-H_2O]^+$ adducts. Chlorophyll a was used to quantify all pigment-related lipids, including chlorophyll, bacteriopheophytin, pyrophaeophorbide a-sterols, carotenoids and their degraded products. For lipid classes without specific standards, such as OLs, a response factor from another class of amino lipids (DGTS) was used. Unknown lipid classes were calibrated using the average response factor of all external standards[34,38].
